# Mechanisms of translation inhibition and suppression of Stress Granule formation by cisplatin

**DOI:** 10.1101/2021.07.20.452628

**Authors:** Paulina Pietras, Anaïs Aulas, Marta M. Fay, Marta Leśniczak, Shawn M Lyons, Witold Szaflarski, Pavel Ivanov

## Abstract

Platinum-based antineoplastic drugs, such as cisplatin, are commonly used to induce tumor cell death. Cisplatin is believed to induce apoptosis as a result of cisplatin-DNA adducts that inhibit DNA and RNA synthesis. Although idea that DNA damage underlines anti-proliferative effects of cisplatin is dominant in cancer research, there is a poor correlation between the degree of the cell sensitivity to cisplatin and the extent of DNA platination. Here, we propose a novel mechanism of cisplatin-mediated cytotoxicity. We show that cisplatin suppresses formation of Stress Granules (SGs), pro-survival RNA granules with multiple roles in cellular metabolism. Mechanistically, cisplatin inhibits cellular translation to promote disassembly of polysomes and aggregation of ribosomal subunits. As SGs are in equilibrium with polysomes, cisplatin-induced shift towards ribosomal aggregation suppresses SG formation and promotes cellular death. Our data also explain nephrotoxic, neurotoxic and ototoxic effects of cisplatin treatment.

## 1. Introduction

Cisplatin [cis-diammine-dichloroplatinum(II)] (CisPt) is a leading antineoplastic platinum-based compound that is widely used to treat roughly 20 distinct tumor types [1]. The clinical benefits of CisPt as an antiproliferative and cytotoxic agent have been recognized for nearly 45 years [2]. While highly effective as a chemotherapeutic agent, CisPt causes a range of side effects including nephrotoxicity, ototoxicity, myelosuppression, gastrotoxicity and allergic reactions [3]. It is assumed, that its closely related analogs carboplatin and oxaliplatin share with CisPt a proposed mechanism of action as DNA-damaging agents, although all compounds demonstrate a difference in the spectrum of toxicities [4]. The cytotoxicity of CisPt is primarily explained by its ability to interact with N7-sites of purine bases in DNA, which promotes formation of both DNA-DNA inter- and intra-strand crosslinks [5] [6]. In turn, such crosslinks distort DNA duplex structures and create CisPt-induced nuclear lesions, the extent of which grossly correlates with extent of cytotoxicity [7]. The CisPt-induced nuclear lesions are proposed to be recognized by different DNA damage proteins or their complexes, which bind to physical distortions on the DNA. They signal then to downstream effectors that promote cascade of signalling events culminating in apoptosis [8].

For decades, it is generally postulated that DNA is a preferential and primary molecular target of CisPt and different types of DNA lesions (monoadducts, inter- and intra-strand crosslinks) trigger DNA damage responses in cells treated with platinum drugs [9]. In agreement with this model, cells deficient in DNA repair are more sensitive to CisPt [10]. However, in enucleated cells CisPt-induced apoptosis occurs independently of DNA damage [11]. Also, less than 1 % of the intracellular CisPt is covalently bound to DNA and there is poor correlation between the sensitivity of cells to the drug and the extent of DNA platination [12]. Moreover, other studies have challenged the DNA-platination model suggesting that CisPt cytotoxicity originates from disrupting RNA processes including induction of ribosomal biogenesis stress [13], inactivation of splicing [14], inhibition of cellular translation [15] [16] and targeting telomeric RNA [17].

Stress granules (SGs) are non-membranous cytoplasmic entities consisting of mRNAs and proteins that form upon cellular exposure to various biotic and abiotic stresses. SGs play critical roles in the Integral Stress Response coordinating multiple cellular processes aimed at promoting cell survival [18]. In cancer cells, SGs confer cytoprotection against chemotherapy, radiotherapy and hostile tumor microenvironment [19]. SGs promote cell survival on multiple levels. SGs block apoptotic pathways by acting as signaling hubs to rewire signaling cascades and act as platforms to re-program cellular translation to conserve energy and redirect that energy to repair stress-induced damage [20]. Additionally, tumor cells promote SG formation to enhance cancer cell fitness and resistance to chemotherapy induced stress thus making SGs potential targets for anti-cancer therapy [19].

Under stress, two major regulatory pathways contribute to SG assembly and modulate protein synthesis by targeting translation initiation [21]. The first pathway targets eukaryotic initiation factor 2 alpha (eIF2α), a component of the eIF2/GTP/tRNAiMet ternary complex that delivers initiator tRNAiMet to the 40S ribosomal subunit. eIF2α is phosphorylated at serine 51 (S51) by one of several stress-activated eIF2α kinases (PKR, PERK, GCN2 and HRI) which inhibits efficient GDP-GTP exchange, prevents the assembly of the ternary complex, and thus inhibits translation initiation (discussed in details in [22]). The second pathway regulates the assembly of the cap-binding eIF4F complex, consisting of eIF4E, eIF4G and eIF4A, controlled by the PI3K-mTOR (mammalian target of rapamycin) kinase cascade. Under optimal conditions, mTOR constitutively phosphorylates its downstream target, eIF4E-binding protein 1 (i.e., eIF4E-BP1 (4E-BP1)) preventing its interaction with eIF4E. Stress-induced inactivation of mTOR leads to the dephosphorylation of 4E-BP1. Dephosphorylated 4E-BP1 prevents the assembly of eIF4F leading to inhibition of translation initiation (reviewed in [23]).

Here, we demonstrate that CisPt affects multiple aspects of mRNA translation by several non-overlapping mechanisms. First, it inhibits translation initiation by promoting 4E-BP1 dephosphorylation and eIF2α phosphorylation. Second, it targets ribosomes and inhibits SG formation in a concentration-and time-dependent manner. CisPt prevents ribosome engagement into translation complexes by inhibiting translation initiation and promoting small ribosomal 40S subunit aggregation in cytosol (CisPt foci). The composition and mechanisms of assembly of CisPt foci are different from canonical SGs [24] [25]. They fail to recruit polyadenylated (poly(A)) mRNAs and lack some SG-associated translation initiation factors. In contrast to SGs, formation of CisPt foci is irreversible and unaffected by pharmacological manipulations of polysomes, translating fraction of ribosomes that form equilibrium with canonical SGs. Formation of CisPt foci sequesters 40S ribosomal subunits and, thus, decreases the number of translating ribosomes, which consequently affecting the formation of SGs. These data provide with alternative explanation on pro-apoptotic effects of CisPt on cancer cells and may explain its cytotoxicity on other non-cancerous cells such as nephrons or inner ear cells where it accumulates under chemotherapy.

## 2. Materials and methods

### 2.1. Cell culture

Human osteosarcoma cells (U2OS, ATCC^®^ HTB-96^™^), mouse embryonic fibroblasts (MEFs) with/without S51A mutation of eIF2*α*, HAP1 cells: (a) parental, (b) eIF2*α* (S51A), (c) ΔHRI, (d) ΔGCN2, (e) ΔPKR, (f) ΔPERK (Horizon Discovery, UK) were grown in Dulbecco’s Modified Eagle Medium with 4.5 g/L D-glucose (DMEM, Gibco) supplemented with 10% fetal bovine serum (Sigma-Aldrich) and Penicillin-Streptomycin cocktail (Sigma-Aldrich).

### 2.2. Antibodies

Anti-G3BP1 (sc-81940; 1:200 dilution for IF), anti-eIF4G (sc-11373; 1:200 dilution for IF, 1:1000 for WB), anti-eIF3b (sc-16377; 1:200 dilution for IF), anti-FXR1 (sc-10554, 1:200 dilution for IF), anti-TIAR (sc-1749; 1:1000 dilution for IF), anti-TIA-1 (sc-1751; 1:1000 dilution for IF), anti-HuR (sc-5261; 1:200 dilution for IF), anti-PABP (sc-32318; 1:100 dilution for IF), anti-p70 S6 kinase (sc-8418, 1:200 dilution for IF) and anti-Rack1 (sc-17754; 1:1000 dilution for WB) were purchased from Santa Cruz Biotechnology (US). Anti-total-eIF2α (#2103, 1:1000 dilution for WB), anti-non-Phospho-4E-BP1 (#4923, 1:1000 dilution for WB), anti-UPF1 (#9435; 1:200 dilution for IF), Caspase 3 total (#9662, 1:1000 dilution for WB) and Cleaved Caspase 3 (#9664, 1:400 dilution for IF) and anti-Rsk2 (#5528; 1:200 dilution for IF) were purchased from Cell Signaling Technology. Anti-Tubulin *α* (66031-1-Ig; 1:1000 dilution for WB), anti-Caprin 1 (15112-1-AP; 1:200 dilution for IF), anti-ABCE1 (14032-1-AP, 1:200 dilution for IF) and PELO (10582-1-AP, 1:200 for IF) were purchased from Protein Technology Group. Anti-ph-eIF2α (Ab32157; 1:1000 dilution for WB) was purchased from Abcam. Anti-Puromycin (MABE343; 1:200 dilution for IF; 1:1000 dilution for WB) was purchased from Millipore. The secondary antibodies for WB, i.e. Peroxidase AffiniPure Donkey Anti-Mouse IgG (cat. 715-035-150) and Peroxidase AffiniPure Donkey Anti-Rabbit IgG (711-035-152) were purchased from Jackson ImmunoResearch. The secondary antibodies for IF included Cy^™^2 AffiniPure Donkey Anti-Mouse IgG (cat. 715-225-150), Cy^™^3 AffiniPure Donkey Anti-Rabbit IgG (711-165-152) and Alexa Fluor^®^ 647 AffiniPure Bovine Anti-Goat IgG (805-605-180) and were purchased from Jackson ImmunoResearch.

### 2.3. Anticancer drugs and chemical compounds

Cisplatin was purchased from BioTang Inc. Cisplatin was prepared directly in DMEM and kept at 4°C. Vinorelbine was purchased from BioTang Inc. Oxaliplatin (commercially available anticancer drug, solution 5 mg/ml) was purchased from Teva Pharmaceuticals, Poland. Carboplatin (commercially available anticancer drug, solution 10 mg/ml) was purchased from Actavis Group PTC, Iceland. Sodium arsenite, puromycin, cycloheximide, and emetine were purchased from Sigma-Aldrich.

### 2.4. Immunofluorescence microscopy

The immunofluorescence technique was done as previously described [26]. Shortly, cells were fixed in 4% paraformaldehyde (Sigma-Aldrich) and permeabilized in cold methanol (−20 °C). Then, cells were incubated with blocking buffer (5% Horse Serum in PBS) for 1 hour. Cells were incubated with primary and secondary antibodies for at least 1 hour each and washed twice with PBS in between incubations. Hoechst 33258 (Sigma-Aldrich) was used together with the secondary antibodies in order to stain the nuclei. Cover slips with cells were mounted in polyvinyl mounting medium. Cells were imaged using an Eclipse E800 Nikon or AxioImager Carl Zeiss microscopes and photographed with either a SPOT CCD or a Pursuit CCD camera (both from Diagnostic Instruments) using the manufacturer’s software. The images were analyzed and merged using Adobe Photoshop CC.

### 2.5. Fluorescence in vitro hybridization (FISH)

10^5^ cells grown on coverslips were fixed in 4% formaldehyde in PBS (15 min) and subsequently permeabilized in 96% cold methanol (15 min). PerfectHyb^™^ Plus Hybridization Buffer (Sigma-Aldrich, H7033) was used to block samples (30 min at 42 °C) and hybridize the probe (synthetic oligo-dT_40_ labeled with cy3 or cy5) for 2h at 42 °C. Then, samples were washed three times with 2xSSC (the first time with pre-wormed and subsequent times with room temperature buffer) and one time with PBS. 0.5 mg/ml UltraPure™ BSA (Ambion, AM2616) was used to block cells and apply primary and secondary antibodies (including Hoechst 33258). Finally, coverslips with cells were washed twice with PBS and mounted in polyvinyl mounting medium.

### 2.6. Western blotting

Cells were grown in 6-well plates until 80% confluence. They were washed with HBSS buffer and solubilized in the lysis buffer (5mM MES, pH 6.2, and 2% SDS), followed by 2 × 2 min sonication at 4 °C. Lysates were denatured in a boiling water and cooled to room temperature. Proteins were precipitated in 60% acetone at -20°C overnight. Lysates were then centrifuged (13500 rpm, 4 °C, 15 min) and supernatant was carefully removed and discarded. Pellets were dissolved in 1x Laemmli loading buffer, proteins were separated in 4-20% SDS-PAGE gels (BioRad) and transferred to nitrocellulose membranes using Trans-Blot^®^ Turbo™ system (BioRad). After 1h blocking in 2% milk in TBS-Tween, membranes were incubated with primary and secondary antibodies for a minimum 1h (membranes were also washed 5x after each type of antibodies). Finally, HRP-conjugated secondary antibodies were detected with SuperSignal West Pico Chemiluminescent Substrate (ThermoScientific) according to the manufacturer instruction.

### 2.7. Quantification of SGs

The percentage of stress granules in a cell population was quantified by manual counting of approximately 700 cells with/without stress granules using Adobe Photoshop CC. Quantification of band intensity in WB technique was done using ImageJ software.

### 2.8. Polysomes profiles

Cells were washed with cold HBSS, scrape-harvested directly into lysis buffer (10 mM HEPES pH 7.5, 125 mM KCl, 5 mM MgCl_2_, 1 mM DTT, 100 μg/mL cycloheximide, 100 μg/mL heparin, 1% NP40 made in DEPC-treated water), supplemented with RNasin Plus inhibitor (Promega) and HALT phosphatase and protease inhibitors (Thermo Scientific). Lysates were rotated at 4°Cfor 15 min, cleared by centrifugation for 10 min at 12,000 *g*, and supernatants loaded on pre-formed 17.5-50% sucrose gradients made in gradient buffer (10 mM HEPES pH 7.5, 125 mM KCl, 5 mM MgCl_2_, 1 mM DTT). Samples were centrifuged in a Beckman SW140 Ti rotor for 2.5h at 35,000 rpm, then eluted using a Brandel bottom-piercing apparatus connected to an ISCO UV monitor, which measured the eluate at OD 254.

### 2.9. Fluorescence recovery after photobleaching (FRAP)

U2OS stably expressing GFP-G3BP1 were plated the day prior the experiment. Cells were stressed as indicated and 30 min before starting the experiment cells were transferred to the FRAP chamber (37 °C, 5 % CO_2_, humidified). 3 frames were collected before bleaching and 20 after, all with an interval of 5 sec in-between. The photobleaching beam was positioned directly over each SG, and laser power were turn to 100 % of the power to perform bleaching.

### 2.10. Ribopuromycylation assay

Ribopuromycylation assay was modified from [27], as described in [28]. In brief, 5 min before fixation, puromycin (Sigma-Aldrich) was added to a final concentration of 5 µg/ml, respectively, and the incubation continued for 5 min. Cells were then lysed subjected to either western blotting or immunofluorescence using anti-puromycin antibody (both techniques as described above). Cells without puromycin treatment were used as negative controls.

## 3. Results

### 3.1. Cisplatin induces formation of SG-like cytoplasmic foci

Recently, we screened a panel of FDA-approved chemotherapy drugs and identified specific compounds that promote SG formation (data not shown). Using the SG-specific marker G3BP1, we tested whether platinum-based drugs such as CisPt, oxaliplatin (OxaPt) and carboplatin (CrbPt) also stimulate formation of canonical SGs as sodium arsenite (SA) and vinorelbine (VRB) in human osteosarcoma U2OS cells (Fig. 1A and Fig. S1A). Indeed, all tested platinum drugs induce formation of G3BP1-positive cytoplasmic foci in 20-40% of cells (Fig. S1B). To characterize these foci, we focused on CisPt as a representative member of platinum drugs. In contrast to SA-and VRB-induced SGs, CisPt-induced foci only contain some of the canonical SG markers, including TIAR and the small ribosomal subunit protein RPS6 (Fig. 1B) but lacking eIF3b, eIF4G (Fig. 1A). Just as SA-induced SGs, large ribosomal subunit protein P0 are not found in CisPt induced foci (Fig. 1D). Using fluorescence in situ hybridization (FISH) to detect polyadenylated mRNAs [25], we failed to identify mRNAs (Fig. 1C) in CisPt-induced foci, in contrast to SA-induced SGs (Fig. 1C-D). In contrast with the failure to efficiently recruit polyadenylated mRNAs to CisPt foci (Fig. 1C), we observe a weak signal for poly (A)-binding protein (PABP) in CisPt-induced foci (Fig. S1D). To determine whether CisPt foci resemble P-bodies (PBs), RNA granules closely related to SGs, we assessed the classical P-body marke Dcp1, a classical marker of PBs (Fig. S1C). In addition to U2OS osteosarcoma cells, CisPt potently induces G3BP1-positive, eIF4g and eIF3b-negative foci in other cancer cell lines (including HeLa (cervix), MCF7 (breast) and A549 (lung)) under similar doses (data not shown).

**Fig. 1.**
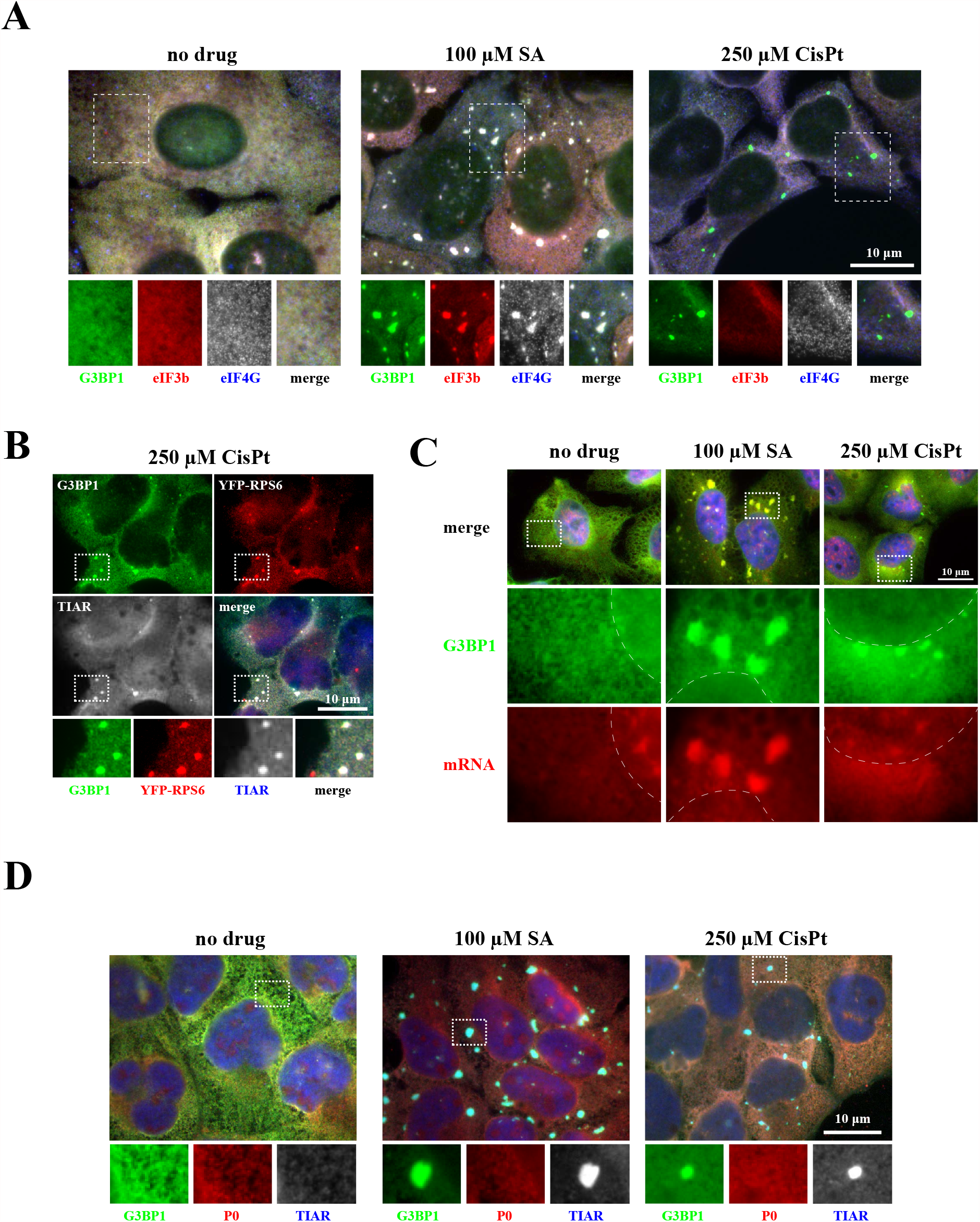
CisPt induces formation of cytoplasmic foci (*A*) U2OS cells were stressed with sodium arsenite (SA, 100 µM) for 1 hour and cisplatin (CisPt, 250 µM) for 4 hours. Unstressed U2OS cells (no drug) were used as control. After drug exposure, cells were fixed and stained for canonical SGs markers: G3BP1 (green), eIF3b (red) and eIF4G (blue, shown as gray). The region in box is enlarged below each image (separated colors in RGB system). The size bar represents 10 µm. (*B*) CisPt-induced granules contain ribosomal protein S6 (red) and TIAR (blue). (*C*) CisPt-induced granules do not contain mRNA. U2OS cells were treated with sodium arsenite (SA, 100 µM) for 1 hour and cisplatin (CisPt, 250 µM) for 4 hours (control cells, untreated, no drug). Cells were fixed and stained for G3BP1 and mRNA using FISH technique (G3BP1 -green -cyanine 2, mRNA -red -cyanine 3 fused with the anti-biotin secondary antibodies; *in situ* hybridization was done using oligo-dT40 probe against polyadenylated mRNA). Nuclei were visualized with Hoechst staining (blue). Boxed region is shown enlarged below each image (each fluorescence channel demonstrated as gray), dotted line represents boundaries of nuclei (*D*) CisPt-induced granules do not contain protein P0 associated with 60S subunit. U2OS cells were stressed with sodium arsenite (SA, 100 µM) for 1 hour and cisplatin (CisPt, 250 µM) for 4 hours (one population of U2OS cells were used as control -no drug). Cells were fixed and stained for three different proteins -G3BP1 (green), P0 (red), TIAR (blue). Boxed region was shown enlarged with colors below each image. The size bar represents 10 µm.

Together, these data show that CisPt-induced foci contain some “canonical” SG components but lack others, most notably polyadenylated mRNAs and early translation initiation factors eIF3b and eIF4G. As recruitment of specific signaling and apoptosis-related molecules into SGs affects stress adaptation and survival of cells (reviewed in [19]), we next examined their localization after CisPt treatment (Fig. S2). p70 S6 kinase, TRAF2 and RSK2 localize to CisPt-induced foci similar to SA-induced SGs suggesting that although their composition is different, signaling molecules still shuttle into CisPt-induced foci similarly to SGs. However, CisPt-induced foci represent some similarities to SGs observed as intense shuttling of G3BP1 in and out of SGs (Fig. 2C).

**Fig. 2.**
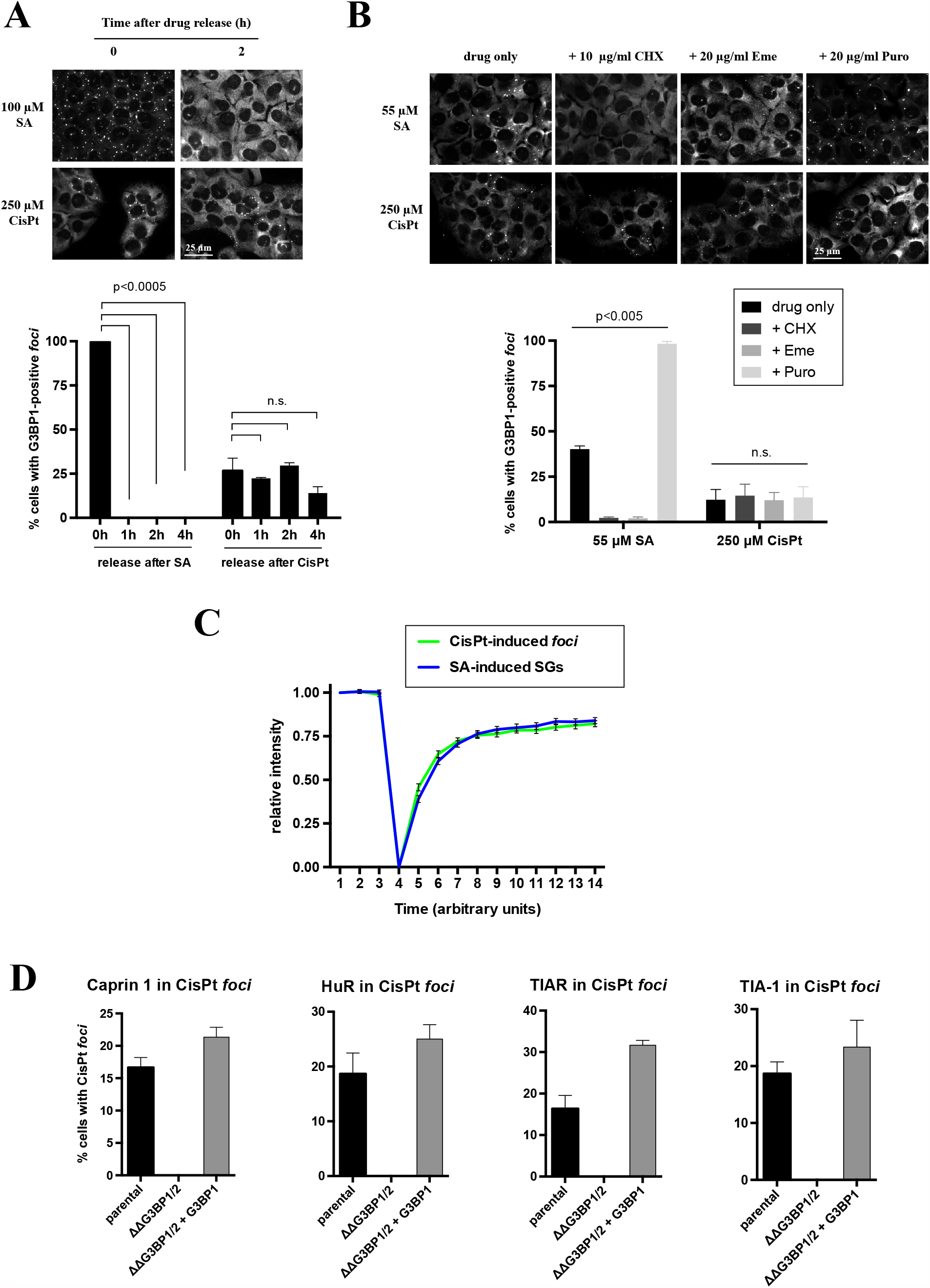
Features of CisPt-granules: dynamics and dependents on G3BP (*A*) Dynamics of CisPt foci after stress relief. U2OS cells were treated either with sodium arsenite (SA, 100 µM) or cisplatin (CisPt, 250 µM) for 1 hour and 4 hours, respectively (negative control not shown). Then drug was removed form media and cells were incubated for additional 1, 2, and 4 hours (control cells were fixed also directly after drug release -indicated as 0h). Cells were fixed and stained for G3BP1 (green) and Hoechst (blue). Data were analyzed using the unpaired Student’s *t*-test, N=3, and demonstrated on the graph. (*B*) Effects of translation inhibitors on CisPt foci formation. Cells were treated either with sodium arsenite (SA, 55 µM) or cisplatin (CisPt, 250 µM) for 1h and 3.5h followed by 1h incubation with cycloheximide (CHX, 10 µg/ml), Emetine (Eme, 20 µg/ml) or Puromycin (Puro, 20 µg/ml). Cells were fixed and stained for G3BP1 (green) and Hoechst (blue). Data were analyzed using the unpaired Student’s *t*-test, N=3, and demonstrated on the graph. (*C*) Quantification of CisPt-granules dynamics using FRAP technique. U2OS stable cell line GFP-G3BP1 was used. 3 frames were collected before bleaching and 20 after, all with an interval of 5 sec in-between. (*D*) G3BP is absolutely required for recruitment of selected SG markers into CisPt foci. Recruitment of into CisPt foci in SG-competent (parentalCaprin1, HuR, TIAR and TIA-1 U2OS, ΔΔG3BP1/2+G3BP1) and SG-incompetent (ΔΔG3BP1/2) U2OS cells.

### 3.2. Cisplatin-induced foci are dynamically distinct from SGs

SGs are dynamic entities that assemble during stress and disassemble upon stress removal [29], and therefore we examined whether CisPt-induced foci dissolve after CisPt is removed from cells. In contrast to the rapid disassembly of SA-induced SGs, CisPt-induced foci are stable two hours after stress removal (Fig. 2A). SGs are in equilibrium with polysomes [29], actively translating fraction of ribosomes, and pharmacological manipulations that affect polysome dynamics also alter SG assembly and disassembly. Cycloheximide (CHX) and emetine (Eme) stall translating ribosomes causing polysome stabilization [25]. This results in the disassembly of SA-induced SGs (Fig. 2B). However, these drugs failed to effect CisPt-induced foci (Fig. 2B, CHX and Eme). Puromycin (Puro) is a translation inhibitor that collapses polysomes by premature termination and promotes SG assembly [25]. Puro treatment enhances the formation of SA-induced SGs but does not influence CisPt-induced foci (Fig. 2B, Puro).

SGs are also dynamic in the movement of components in and out of the granule [30]. We monitored the residence time of GFP-tagged G3BP1 using Fluorescence Recovery After Photobleaching (FRAP) in SA-induced SGs and CisPt-induced foci. In these experiments, the behavior of G3BP1 was similar in both SGs and CisPt-induced foci (>90% recovery of the bleached signal occurred within 10 s) suggesting that G3BP1 rapidly shuttling in and out of CisPt-induced foci (Fig. 2C). Collectively, these data indicate that CisPt-induced are not SGs, although they share some components with canonical SGs (Figures 1 and 2).

G3BP is a protein critical for SG formation under many stresses [31]. We tested whether G3BP is also required for the assembly of CisPt-induced foci using U2OS cell line with genetic knockout of both G3BP proteins (ΔΔG3BP1/2) [31]. Recruitment of SG markers Caprin 1, HuR, TIAR and TIA-1 into CisPt foci is completely abolished when compared to parental U2OS cells (Fig. 2D). SG recruitment defects are efficiently rescued by expression of G3BP1 (Fig. 2D, ΔΔG3BP1/2+G3BP1). This suggests that G3BP is required for CisPt foci formation similarly as for canonical SGs.

### 3.3. Cisplatin-induced foci are formed as a result of translation repression

Canonical SGs form when translation initiation is inhibited. To determine whether CisPt-induced foci are connected to translation initiation inhibition, we first examined whether CisPt treatment alters cellular translation using polysome profiling, a fractionation method of grossly assessing the overall translational state of cells.

Polysome profiling indicates that CisPt promotes disassembly of polysomes and accumulation of monosomes and ribosomal subunits, although less potently than SA that was used as a control (Fig. 3A).

**Fig. 3.**
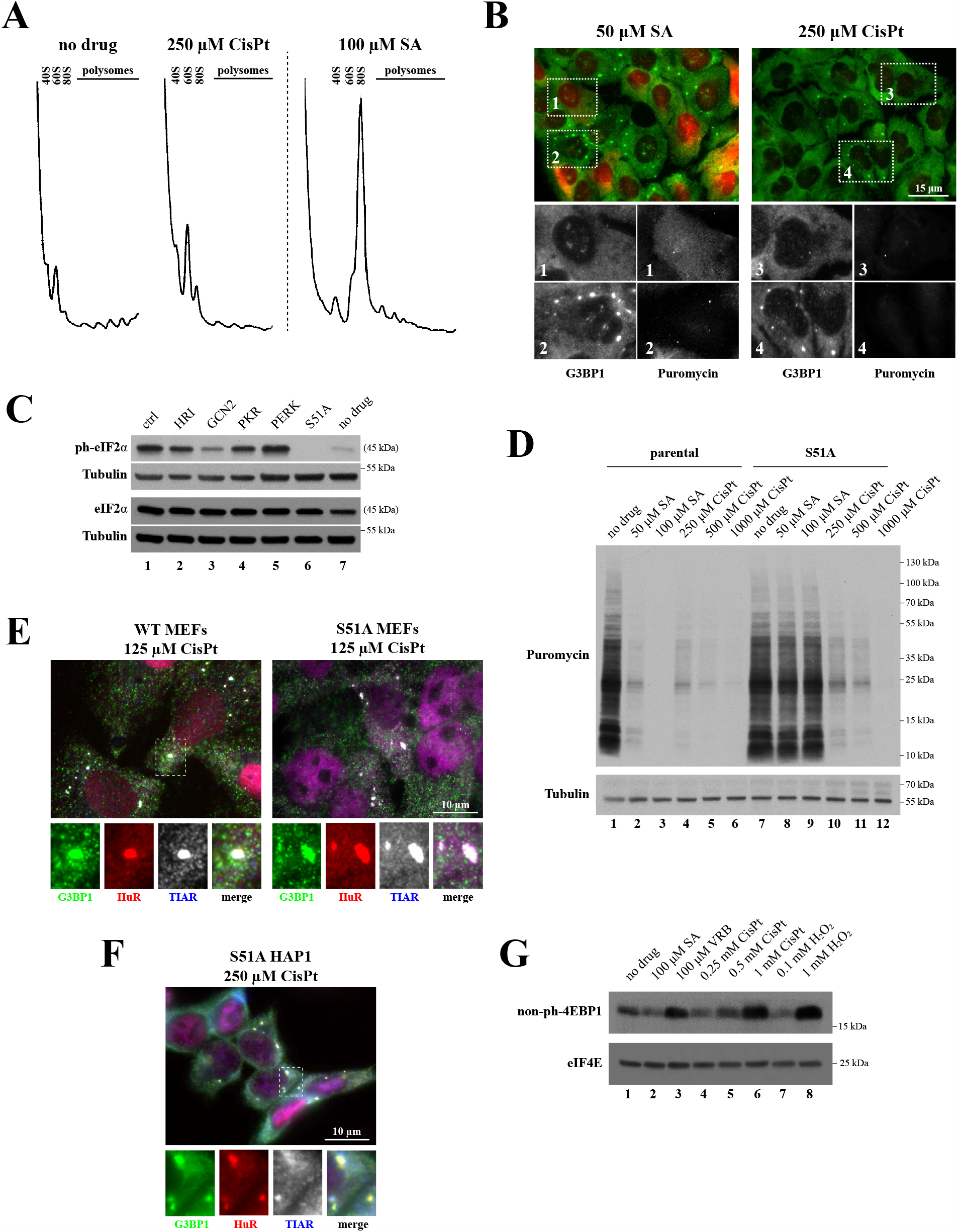
CisPt inhibits translation by promoting eIF2*α* phosphorylation and inhibition of mTOR (*A*) Polysome profiles obtained from U2OS cells treated with cisplatin (CisPt, 250 µM) or sodium arsenite (SA, 100 µM) for 4 hours and 1 hours, respectively; as control, polysomes were isolated from untreated control cells (no drug). (*B*) Detection of translation activity based on immunofluorescence technique. U2OS cells were treated with sodium arsentite (SA, 50 µM) or cisplatin (CisPt, 250 µM), fixed and stained with G3BP1 (green) to detect CisPt foci and anti-puromycin to monitor translation (red, shown as grey in boxed sub-image). The size bar represents 10 µm. (*C*) Effect of CisPt on eIF2*α* phosphorylation. Parental HAP1 (ctrl), HAP1 variants with eIF2*α* kinase knockout genes (HRI, GCN2, PKR and PERK) or with eIF2*α* S51A mutation (S51A) were treated with CisPt (250 µM, 4 hours). Untreated parental HAP1 cells (no drug) were used as controls. Lysates from treated and control cells were analyzed by western blotting using phosphor-eIF2*α* antibody (ph-eIF2*α*) Total eIF2*α* (eIF2*α* on the Fig.) and tubulin (Tubulin on the Fig.) were used as loading controls. (*D*) Detection of translation activity in two populations of HAP1 cells (parental and S51A) treated with sodium arsenite (50 µM, 100 µM, SA), cisplatin (250 µM, 500 µM, 1000 µM, CisPt). No treated control (No drug) was used as control. U2OS cells were subjected to RiboPuromycylation to compare levels of basal translation. An anti-puromycin antibody (Puro) was used to visualize *de novo* synthesized proteins. Tubulin is a loading control. A representative image is shown (*n*=3). (*E-F*) Formation of CisPt foci is independent of eIF2*α* phosphorylation. E: formation of CisPt foci in WT and S51 mouse embryonic fibroblasts (MEFs). F: formation of CisPt foci in S51 HAP1 cells. G3BP1, HuR and TIAR were used as markers. (*G*) CisPt promotes eIF4EBP1 dephosphorylation and its binding to eIF4E. U2OS cells were stressed with the following chemicals: SA (100 µM, 1h), VRB (100 µM, 1h), CisPt (0.25 mM, 4h), CisPt (0.5 mM, 4h), CisPt (1 mM, 4h), H_2_O_2_ (0.1 mM, 1h), H_2_O_2_ (1 mM, 1h). Untreated U2OS cells (no drug) were used as controls. Cells were lysed and subjected to m^7^GTP-sepaharose pull down to isolate cap-bound complexes. Precipitated (m7GTP) fractions were subjected to western blotting and probed against eIF4E and non-phosphorylated 4E-BP1 (non-ph-4EBP1).

Ribopuromycylation, a technique that directly assesses translation activity in cells, demonstrates that CisPt potently inhibits translation in both cells that assemble (Fig. 3B, box 4) and do not assemble (Fig. 3B, box 3) CisPt-induced foci. This is in contrast to SA, where translation inhibition and SG assembly are coupled (Fig. 3B, compare boxes 1 and 2).

Two main pathways regulate translation in response to stress, both targeting translation initiation: 1) control of initiator tRNA delivery to the ribosome by phosphorylation/dephosphorylation of eIF2α, and 2) mTOR-regulated binding of eIF4E-BPs to cap-binding protein eIF4E. In HAP1 cells [24], CisPt triggers robust eIF2α phosphorylation (ph-eIF2α, compare lanes 1 (ctrl) and 7 (no drug), Fig. 3C and S3) but does not affect eIF2α protein levels (Fig. 3C, lower panel). CisPt-induced eIF2α phosphorylation is mediated by GCN2 kinase as GCN2 knockout cells (Fig. 3C, lane 3) but no other eIF2α kinases show decreased levels of ph-eIF2α (Fig. 3C, lanes 2-5). HAP1 cells bearing a non-phosphorylatable eIF2α mutant with Ala to Ser substitution at the position 51 (S51A) were used as control (Fig. 3C, lane 6). Further, puromycin labeling demonstrates that CisPt inhibits translation in both WT (Fig. 3D and S3, compare lanes 4-6 with lane 1 (no treatment)) and eIF2α-S51A HAP1 cells (Fig. 3D, compare lanes 10-12 with lane 7 (no treatment)). This is in contrast to SA, which inhibits translation only in WT but not S51A HAP1 cells (Fig. 3D, lanes 1-3 and 7-9). This indicates that unlike SA, CisPt-induced translation repression is not entirely dependent on eIF2α.

To determine whether eIF2α is required for CisPt-induced foci assembly, we treated eIF2α-S51A mutant mouse embryonic fibroblasts (MEFs, Fig. 3E) and eIF2α-S51A HAP1 cells (Fig. 3F) with CisPt. In both cases, CisPt-induced foci are formed suggesting that foci formation is not exclusively dependent upon eIF2α phosphorylation. As it is seen in Fig. 3G (and S3), CisPt promotes dephosphorylation of 4E-BP1. This is similar to the previously reported effect of hydrogen peroxide (H_2_O_2_, [32]) and vinorelbine (VRB, [26]) on 4E-BP1, which were used as controls. Furthermore, nitric oxide-induced inhibition of protein synthesis results from both phosphorylation of eIF2α and displacement of the eIF4F complex as a result of 4EBP dephosphorylation [33]. In contrast, and in agreement with previous observations, SA does not cause 4E-BP dephosphorylation. Thus, CisPt triggers both dephosphorylation of 4EBP and phosphorylation of eIF2α to inhibit translation.

### 3.4. Cisplatin suppresses formation of Stress Granules

Our data suggest that CisPt promotes formation of 40S-containing cytoplasmic foci by inhibition of cellular translation (Figures 1 and 2). As 40S ribosomal subunits are core constitutes of SGs, we hypothesize that CisPt-induced accumulation of 40S subunits into these foci limits pool of ribosomes available for protein biosynthesis. Moreover, since polysomes are in equilibrium with SGs, we predicted that by decreasing the pool of actively translated ribosomes, CisPt will negatively affect formation of SGs. We pretreated cells with different concentrations of CisPt and then followed by a treatment with SA (Fig. 4). As can be judged by the recruitment of SG markers eIF4G and eIF3b, CisPt pre-treatment with low amounts of CisPt (10-50 µM, 24h) causes dose-dependent, statistically significant, decrease of SG-positive cells (Fig. 4A). U2OS cells treated with higher concentrations of CisPt (250 µM, Fig. 4B) readily demonstrate significantly reduced SG formation at shorter times (1 to 3 hours). Thus, treatment with CisPt promotes formation of CisPt foci that reduce abilities of cell to promote SG formation in response to stress.

**Fig. 4.**
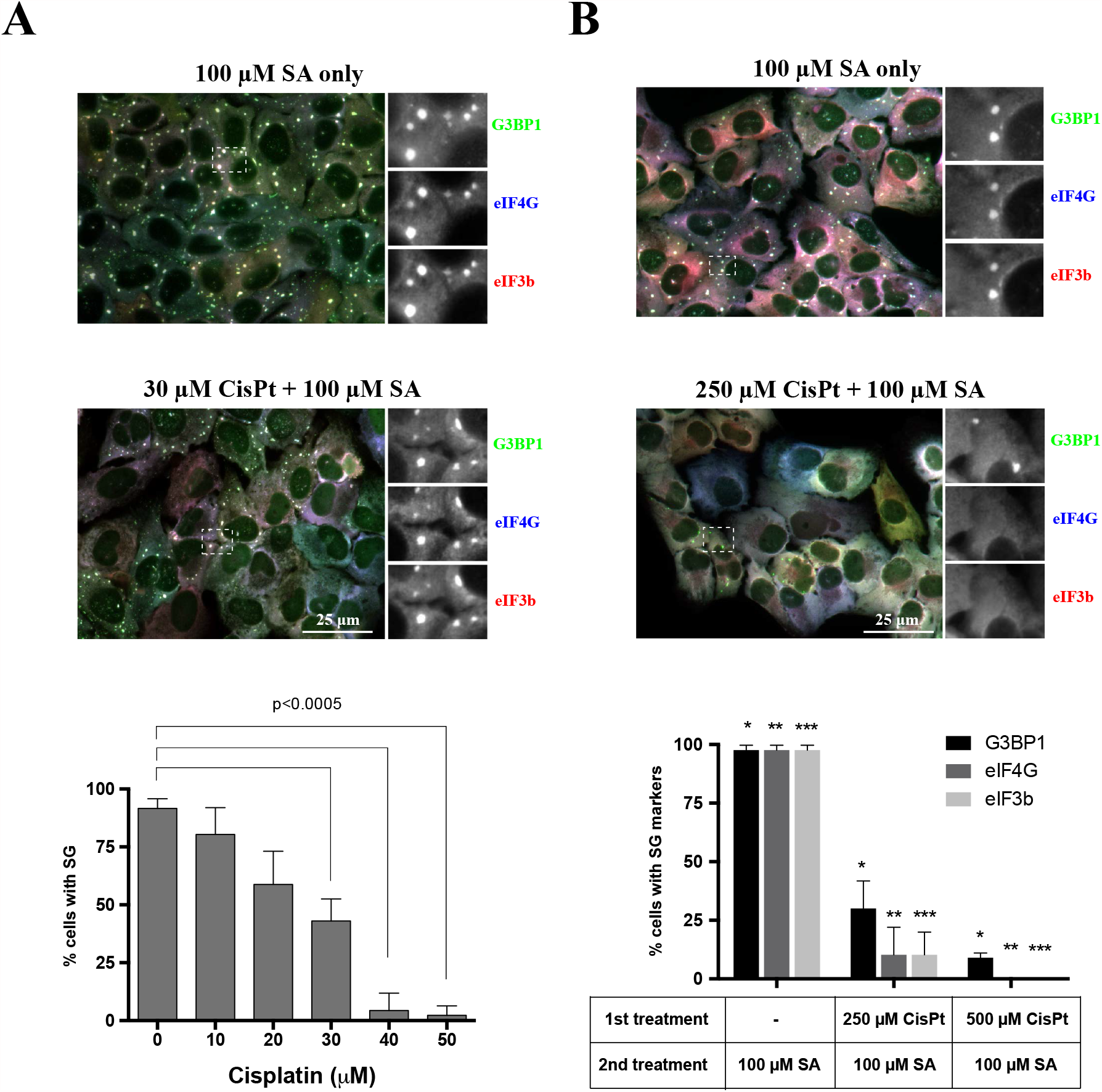
CisPt suppresses SG formation (*A*) The formation of SA-induced stress granules was tested in U2OS cells in two populations of U2OS cells. Control population (untreated) and previously pretreated with increasing amount of cisplatin (0-50 µM) for 24 hours. The cells from both populations were stressed with 100 µM sodium arsenite for 1 hour. The upper image demonstrates population of U2OS cells stressed only with sodium arsenite (SA only, 100 µM), the lower image shows population of U2OS cells pretreated with cisplatin for 24 hours and treated with sodium arsenite for 1 hour (30 µM CisPt + 100 µM SA); representative image. The cells were stained for canonical stress granules markers: G3BP1 (green), eIF4G (blue) and eIF3b (red). The main image was merged (RGB system). Boxed region was shown enlarged in grey corresponding to specific fluorescence channel as indicated. Data were analyzed using the unpaired Student’s *t*-test, N=3, and demonstrated on the graph. (*B*) Control population (untreated) and previously pretreated with CisPt (no drug, 250 µM, 500 µM) for 3 hours. The cells from both populations were stressed with 100 µM sodium arsenite for 1 hour. The upper image demonstrates population of U2OS cells stressed only with sodium arsenite (SA only, 100 µM), the lower image shows population of U2OS cells pretreated with cisplatin for 3 hours and treated with sodium arsenite for 1 hour (250 µM CisPt + 100 µM SA). The cells were stained for canonical stress granules markers: G3BP1 (green), eIF4G (blue) and eIF3b (red). The main image was merged (RGB system). Boxed region was shown enlarged in grey corresponding to specific fluorescence channel as indicated. Data were analyzed using the unpaired Student’s *t*-test, N=3, and demonstrated on the graph. The size bar represents 10 µm.

## 4. Discussion

Cisplatin plays a key role in cancer chemotherapy where it is highly effective against a variety of solid tumors [1, 3]. Historically, DNA is generally considered as a major biological target of CisPt. Upon entering the cell, CisPt is activated through a serious of spontaneous aquation reactions resulting in the generation of a powerful electrophile [34, 35]. The monoaquated form represents as a highly reactive species, which formation is regulated by the interaction with a number of intracellular nucleophiles. These endogenous nucleophiles such as proteins, glutathione or methionine contribute to the intracellular inactivation of CisPt thus modulating its bioactivity.

This simple model where DNA damage underlines CisPt cytotoxicity is challenged by other studies. They suggest that CisPt cytotoxicity originates from multiple sources besides DNA damage-mediated [36], e.g. by targeting RNA metabolism by interference with telomerase functions [17], or inhibition of protein synthesis, transcription and splicing [37]. Moreover, experiments on enucleated cells demonstrated that CisPt-induced cytotoxicity does not involve DNA damage [11]. In agreement with it, only limited amount of intracellular CisPt is covalently bound to DNA, and there is no linear correlation between the extent of DNA platination and its toxicity to cells [12]. Thus, the ability of CisPt to induce nuclear DNA damage *per se* is not sufficient to explain its high degree of effectiveness on highly proliferative cancer cells nor the cytotoxic effects exerted on normal, post-mitotic tissues.

Our analysis reveals new mechanisms of CisPt-mediated cytotoxicity. Our data suggest that CisPt strongly affects RNA metabolism by modulation of common stress responses acting on post-transcriptional level. We show that CisPt potently inhibits cellular translation (Fig. 3B, D). This CisPt-mediated inhibition of protein synthesis is mediated by inactivation of mTOR leading to dephosphorylation of 4EBP1 (Fig. 3G) and by phosphorylation of eIF2*α* via activation of the GCN2 kinase (Fig. 3C). Both mTOR inactivation and phosphorylation of eIF2*α* lead to the inhibition of translation initiation and partial reduction of polysomes (Fig. 3A). Activation of both pathways is likely to be a consequence of mTOR and GCN2 sensing reactive oxygen species induced by CisPt-mediated damage of mitochondrial species [38] rather than by direct interaction with drug.

Inhibition of translation initiation is commonly coupled with formation of SGs [21]. SGs form in response to various extra-and intra-cellular insults and aim on stress adaptation [39]. SGs can promote viability by several mechanisms, which serve to conserve and redirect cellular energy towards pro-survival strategies. Several chemotherapy agents have been previously reported to promote SG formation. In contrast to these drugs, CisPt induces formation of unique cytoplasmic foci that are distinct from SGs, although share with them some canonical components such as 40S ribosomal subunits, markers G3BP1, TIAR or PABP (Fig. 1 and Fig. S1) as well as some signalling molecules (Fig. S2). CisPt foci are also different from P bodies (Fig. S1C), other well-known cytoplasmic RNA granules. Albeit the presence of 40S subunits, CisPt foci lack poly(A) mRNAs (Fig. 1C) that can explain the absence of initiation factors eIF3b and eIF4G (Fig. 1A). It is important to note that although CisPt promotes phosphorylation of eIF2*α*, it promotes CisPt formation in phospho-eIF2*α*-independent manner (Fig. 3E-F). The protein composition of SGs may also be important in predicting the aggressiveness of cancer in patients [40].

Another striking difference of CisPt foci to SGs is that their formation is irreversible (Figures 2A-B). While SGs are quickly dissolving after stress relief, CisPt foci are static and long lived after drug removal (Fig. 2A). In agreement with static nature of CisPt foci, manipulations with polysomes, the fraction of ribosomes that are in dynamic equilibrium with SGs, do not affect formation of these foci (Fig. 2B). In the same time, formation of CisPt foci and SGs is absolutely dependent on the activities of G3BP1, which dynamically associates with 40S subunits and promotes SG condensation. G3BP1 shuttles on and off CisPt foci with kinetics similar to observed with SGs (Fig. 2C), and regulates recruitment of other SG markers into CisPt foci (Fig. 2D). All these data suggest that CisPt foci are both distinct from and related to SGs in terms of their composition and molecular mechanisms of their assembly.

Another possible mechanism of CisPt foci formation is its ability to bind ribosomes directly. The study by the Polikanov laboratory demonstrates that CisPt directly binds to ribosomes and modifies their functional centres such as the mRNA-channel and the GTPase center [41]. By binding to these centres, CisPt interferes with mRNA-ribosome interactions resulting in impaired mRNA translocation and inhibition of protein synthesis. If mechanisms of Cis-Pt binding to ribosomes are conserved between archaea and higher eukaryotes, we propose that CisPt also bind mammalian ribosomes and/or their subunits. By binding to the ribosomes, CisPt inactivates them in a manner that promotes accumulation of 40S subunits into CisPt foci. As 40S subunits are core components of SGs, we predicted that pre-treatment of cells with CisPt would limit available pool of 40S subunits and suppress SGs formation. In agreement with such prediction, incubation of cells with CisPt directly impact their ability to assemble SGs in both time-and concentration-dependent manners (Fig. 4A-B).

The finding that CisPt inhibits SG formation also uncovers new mechanism of CisPt cytotoxicity. As SGs are pro-survival, suppression of their formation contributes to cell death, especially under stress conditions. We propose that in rapidly proliferating cancer cells, suppression of SGs also contribute to CisPt-mediated cell death together with other mechanisms such as DNA damage. However, as CisPt also accumulates in specific cells (nephrons, inner ear cells), which are not cancerous, inhibition of SG formation and protein synthesis may be dominant mechanisms underlying CisPt cytotoxicity.

## CRediT authorship contribution statement

Conceived and designed the experiments: Anaïs Aulas, Marta M. Fay, Marta Leśniczak, Shawn M. Lyons; Analysis of the data and preparation of the draft manuscript: Witold Szaflarski; Wrote the paper: Pavel Ivanov

## Figures and legends

**Fig. S1**. Platin-based drugs induce cytoplasmatic granules formation.

(*A*) Formation of cytoplasmic granules. U2OS cells were stressed with sodium oxaliplatin (OxaPt, 2 mM), carboplatin (CrbPt, 10 mM) and vinorelbine (VRB, 150 µM) for 4 hours (oxaliplatin and carboplatin) and 1 hour (vinorelbine). Unstressed U2OS cells were used as control (no drug; shown in Fig. 1A). After treatment, cells were fixed and stained for stress granules markers: G3BP1 (green), eIF3b (red) and eIF4G (blue, shown as grey). Boxed region is shown enlarged with colors separated below each image. The size bar represents 10 µm.

(*B*) Quantification of cytoplasmatic granules-positive U2OS cells (as in Fig. 1A and Fig. S1A). Data were analyzed using unpaired Student’s *t*-test, N=3.

(*C*) Detection of P-body marker Dcp1 in U2OS cells stressed with cisplatin (CisPt, 250 µM). Control, unstressed, population of U2OS cells was used as control (no drug). After treatment, cells were fixed and stained for G3BP1 (green), Dcp1 (red) and Hoechst (blue). Boxed region is shown enlarged with colors separated below each image; all colors (RGB) are merged in the main image. The size bar represents 10 µM.

(*D*) Detection of typical stress granules markers in cisplatin-stressed U2OS cells. One population of U2OS cells were used as unstressed control (no drug). Cells were stressed with sodium acetate (SA, 100 µM) or cisplatin (CisPt, 250 µM), for 1 hour and 4 hours, respectively. Then, cells were fixed and stained for G3BP1 (green), PABP (red) and TIAR (blue). All channels were demonstrated in grey in box region. The size bar represents 10 µm.

**Fig. S2**. Determination of signaling molecules content in SA-and CisPt-induced SGs. The formation of SGs upon SA and CisPt treatment was checked in U2OS cells. Rsk2, TRAF2 and p70 S6 proteins were detected in SGs using standard immunofluorescence. As a marker of SGs, TIAR or G3BP1 was used. The size bar represents 10 µm.

**Fig. S3**. Uncropped images of gels as related to Figures 3C, 3D and 3G

