## Supplementary figures and images for "Mechanisms of translation inhibition and suppression of Stress Granule formation by cisplatin"

### Supplemental figures

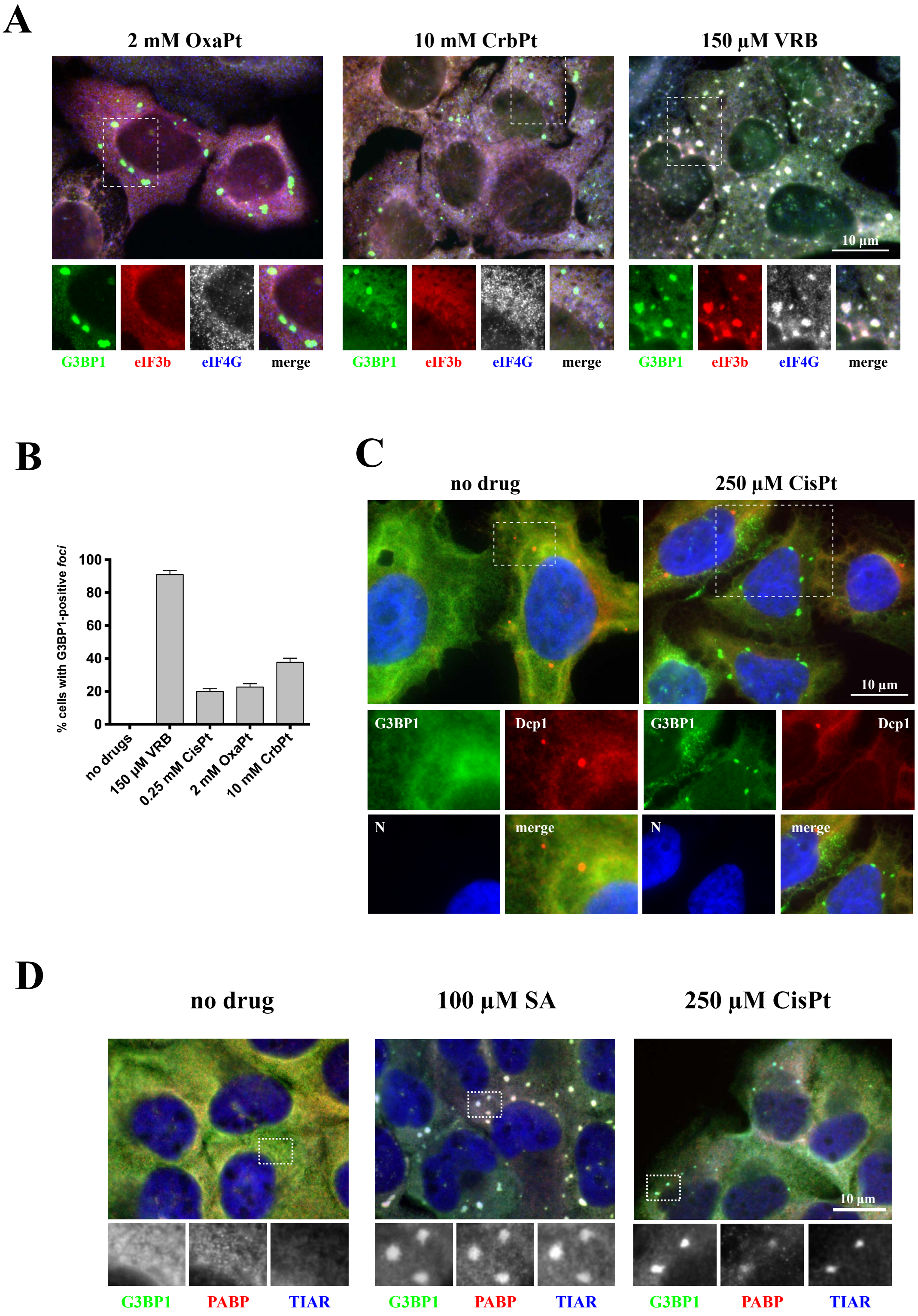

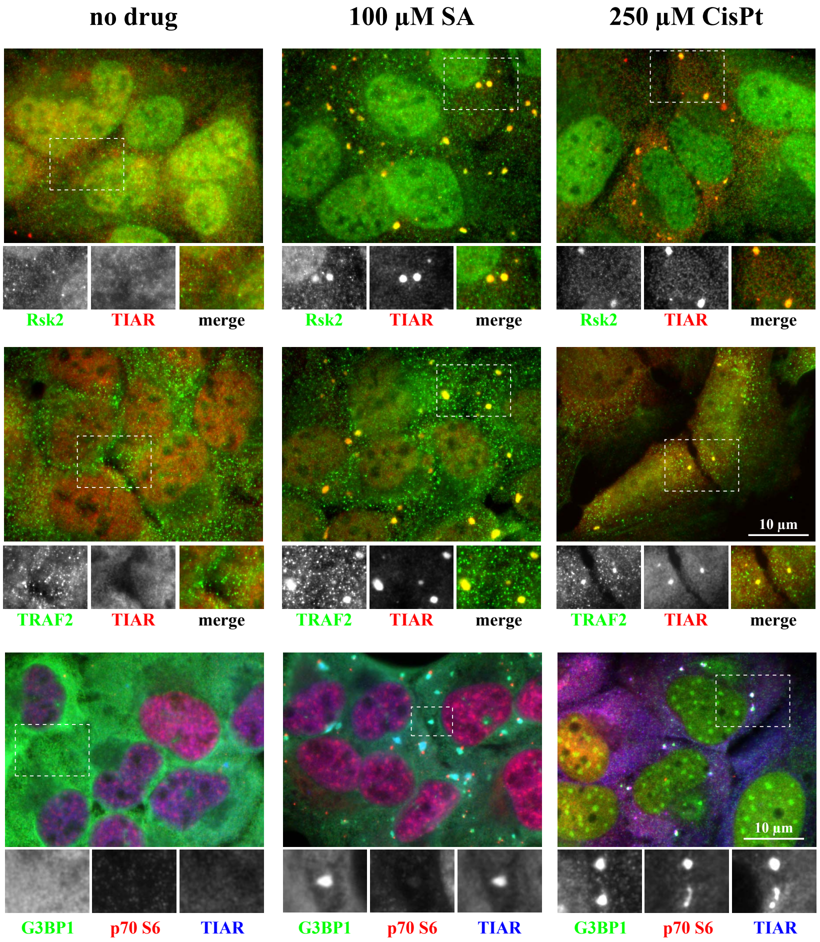

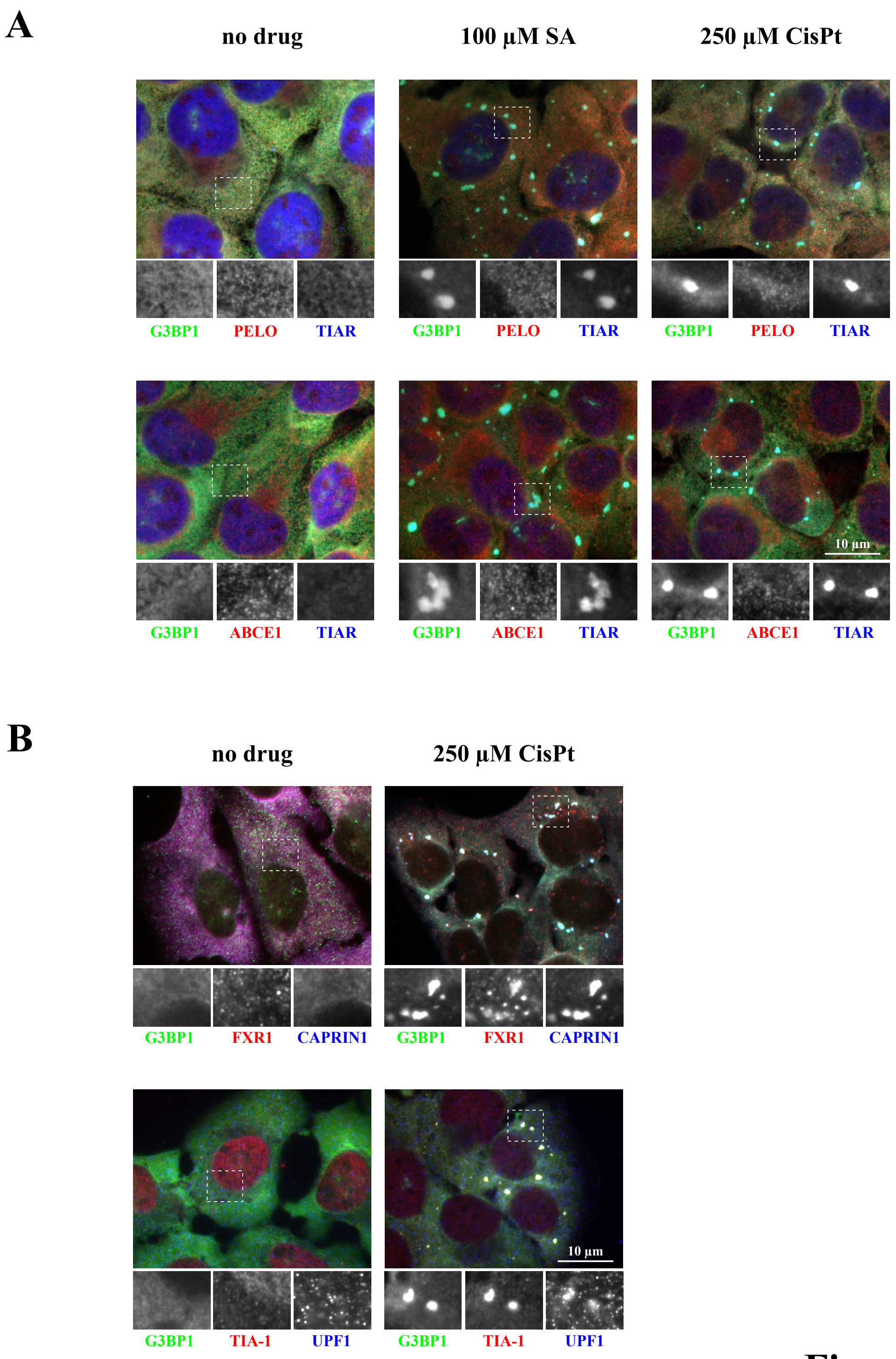

#### Figure 3C

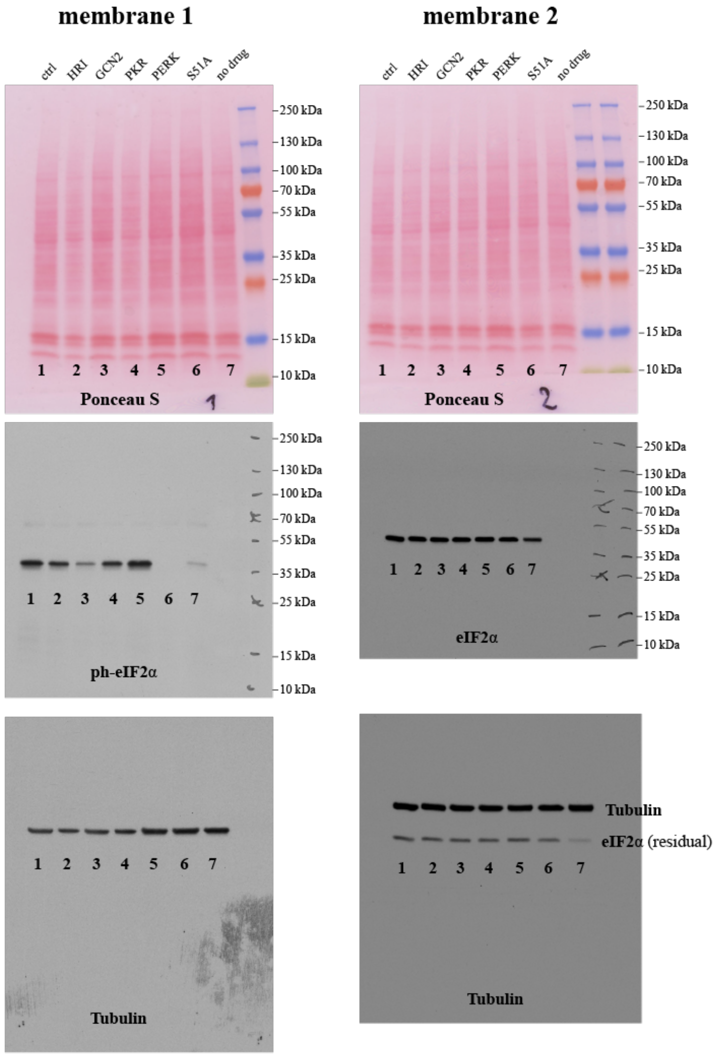


#### Figure 3D

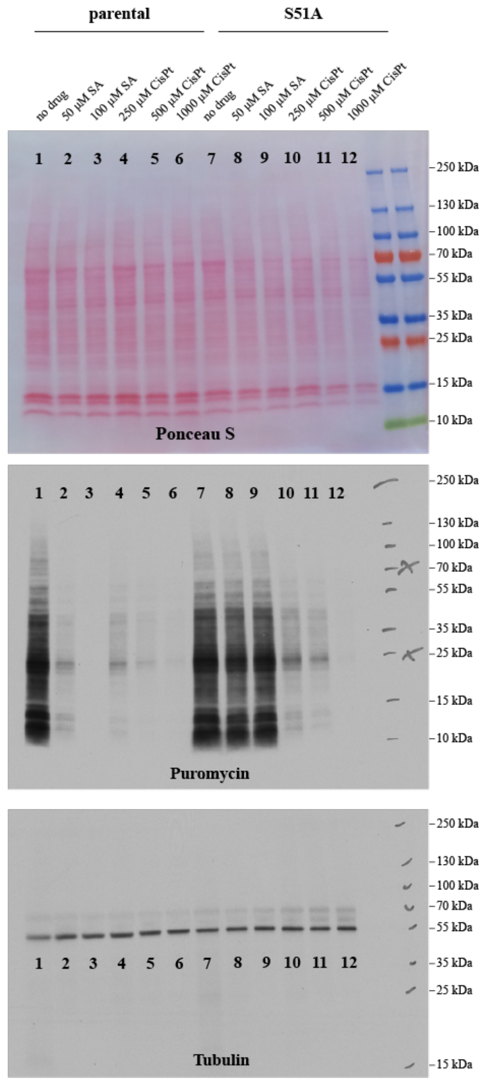


#### Figure 3G

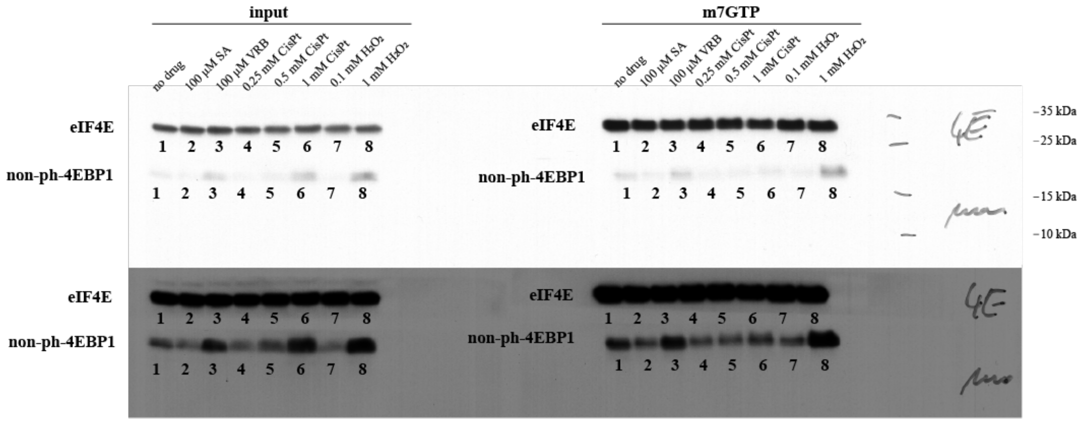


#### Figure S4
